# Gene expression profiles of inflammatory breast cancer reveal high heterogeneity across the epithelial-hybrid-mesenchymal spectrum

**DOI:** 10.1101/2020.08.27.267609

**Authors:** Priyanka Chakraborty, Jason T George, Wendy A Woodward, Herbert Levine, Mohit Kumar Jolly

**Affiliations:** Centre for BioSystems Science and Engineering, Indian Institute of Science, Bangalore 560012, India; Center for Theoretical Biological Physics, Rice University, Houston, TX 77005, USA; Medical Scientist Training Program, Baylor College of Medicine, Houston, TX 77005, USA; Department of Radiation Oncology, The University of Texas MD Anderson Cancer Center, Houston, TX, USA; MD Anderson Morgan Welch Inflammatory Breast Cancer Research Program and Clinic, The University of Texas MD Anderson Cancer Center, Houston, TX, USA; Departments of Physics and Bioengineering, Northeastern University, Boston, MA 02115, USA

**Keywords:** IBC, Gene expression signature, tumor heterogeneity, hybrid epithelial/mesenchymal

## Abstract

Inflammatory breast cancer (IBC) is a highly aggressive breast cancer that metastasizes largely via tumor emboli, and has a 5-year survival rate of less than 30%. No unique genomic signature has yet been identified for IBC nor has any specific molecular therapeutic been developed to manage the disease. Thus, identifying gene expression signatures specific to IBC remains crucial. Here, we compare various gene lists that have been proposed as molecular footprints of IBC using different clinical samples as training and validation sets and using independent training algorithms, and determine their accuracy in identifying IBC samples in three independent datasets. We show that these gene lists have little to no mutual overlap, and have limited predictive accuracy in identifying IBC samples. Despite this inconsistency, single-sample gene set enrichment analysis (ssGSEA) of IBC samples correlate with their position on the epithelial-hybrid-mesenchymal spectrum. This positioning, together with ssGSEA scores, improves the accuracy of IBC identification across the three independent datasets. Finally, we observed that IBC samples robustly displayed a higher coefficient of variation in terms of EMT scores, as compared to non-IBC samples. Pending verification that this patient-to-patient variability extends to intratumor heterogeneity within a single patient, these results suggest that higher heterogeneity along the epithelial-hybrid-mesenchymal spectrum can be regarded to be a hallmark of IBC and a possibly useful biomarker.

## Introduction

Inflammatory breast cancer (IBC) is a rare (2-4% of breast cancer cases) but highly aggressive, locally advanced breast cancer with extremely poor prognosis and a 5-year survival rate of less than 30% [1]. At diagnosis, most IBC patients exhibit signs of lymph node metastasis; approximately 30% of patients have distant metastases as compared to 5% of patients in non-IBC breast cancers [2]. IBCs are often highly angiogenic and invasive, of high histological grade, and cause 7-10% of all breast cancer-associated deaths [1]. Histologically, IBC cells often do not present as a dominant mass, rather being diffused in clusters throughout the breast and skin, thereby leading to many false-negative imaging findings [3]. Decoding unique molecular underpinnings of this deadly disease remains an unmet clinical need.

The presence of tumor emboli in dermal-lymphatic vessels is a pathological hallmark of IBC. Consistently, IBC patients have a higher frequency and larger average size of clusters of circulating tumor cells (CTCs) as compared to non-IBC patients [4]. These clusters have a strong association with poor survival [4], reminiscent of the disproportionately high metastatic fitness of CTC clusters [5]. Aside from this major difference, no unique genomic signature has yet been conclusively identified for IBC, suggesting that other factors may be more important than genetic events for IBC: phenotypic and/or epigenetic heterogeneity, interaction of malignant cells within emboli or with cells from the tumor microenvironment [1,3]. Because of these uncertainties, no specific molecular therapeutic approaches have been yet proposed to manage IBC.

Extensive efforts have been undertaken to identify unique gene expression signatures of IBC. In 2013, the IBC World Consortium identified a 79-gene signature that focused on the inhibition of TGFβ signaling as molecular footprint of IBC [6]; this was consistent with experimental observations that weakened TGFβ signaling promoted collective cell invasion, a hallmark of IBC, while a strong activation of TGFβ signaling promotes individual invasion consistent with a full-blown EMT [7–9]. However, later, these differences were found to arise due to difference in incidence of HER2-positive subtype in IBC vs. non-IBC samples used [1]. Further efforts involving micro-dissected tumors led to identification of 132-gene signature associated with poor outcome [10], but this signature was seen in approximately 25% of breast cancer samples in TCGA which in fact has very few IBC samples, thus highlighting the limited ability of this signature in identifying IBC [1]. The only other study using micro-dissected tumors identified differences in gene expression in the stroma, instead of the tumor cells, in IBC vs. non-IBC cases [11]. Thus, a comprehensive gene expression signature of IBC remains to be identified.

Here, we compare the utility of multiple proposed IBC gene expression signatures by their ability to identify IBC vs. non-IBC cases. Our results here reveal shortcomings in the consistency and predictive utility of these signatures. We subsequently show that single-sample gene set enrichment analysis (ssGSEA) of IBC samples correlate with their EMT scores as calculated via three independent metrics that quantify the spectrum of epithelial-hybrid-mesenchymal phenotypes. Next, we show that while mean EMT scores of IBC samples were not consistently high or low when compared with corresponding non-IBC samples, IBC samples robustly displayed a comparatively higher coefficient of variation in terms of their EMT score. These results suggest that higher heterogeneity along epithelial-hybrid mesenchymal spectrum can be regarded a hallmark of IBC.

## Results

### Available IBC gene signatures do not distinguish robustly between IBC and non-IBC

First, we collated the four available gene lists identified to be unique to IBC. These four gene signatures showed high accuracy in classifying IBC samples from non-IBC (nIBC) samples in their respective studies. Each signature was comprised of a different number of genes/probes – 132, 78, 50 and 109 (denoted as 132 GES, 78 GES, 50 GES and 109 GES henceforth) [6,10,12,13]. The 109 GES signature consists of 109 different probe sets which uniquely mapped to 90 genes; in all other gene signatures, all probe sets mapped to an equal number of genes. These gene signatures were identified via distinct data-driven algorithms, each utilizing a single dataset and variable number of samples in the respective training and validation sets. These signatures exhibited varied accuracy in identifying IBC cases (**Fig 1A**). Investigating the intersection of common genes amongst each signature revealed minimal or no overlap. Aside from one common gene identified by 132 GES and 50 GES, all other signature elements were unique (**Fig 1B**).

**Fig 1:**
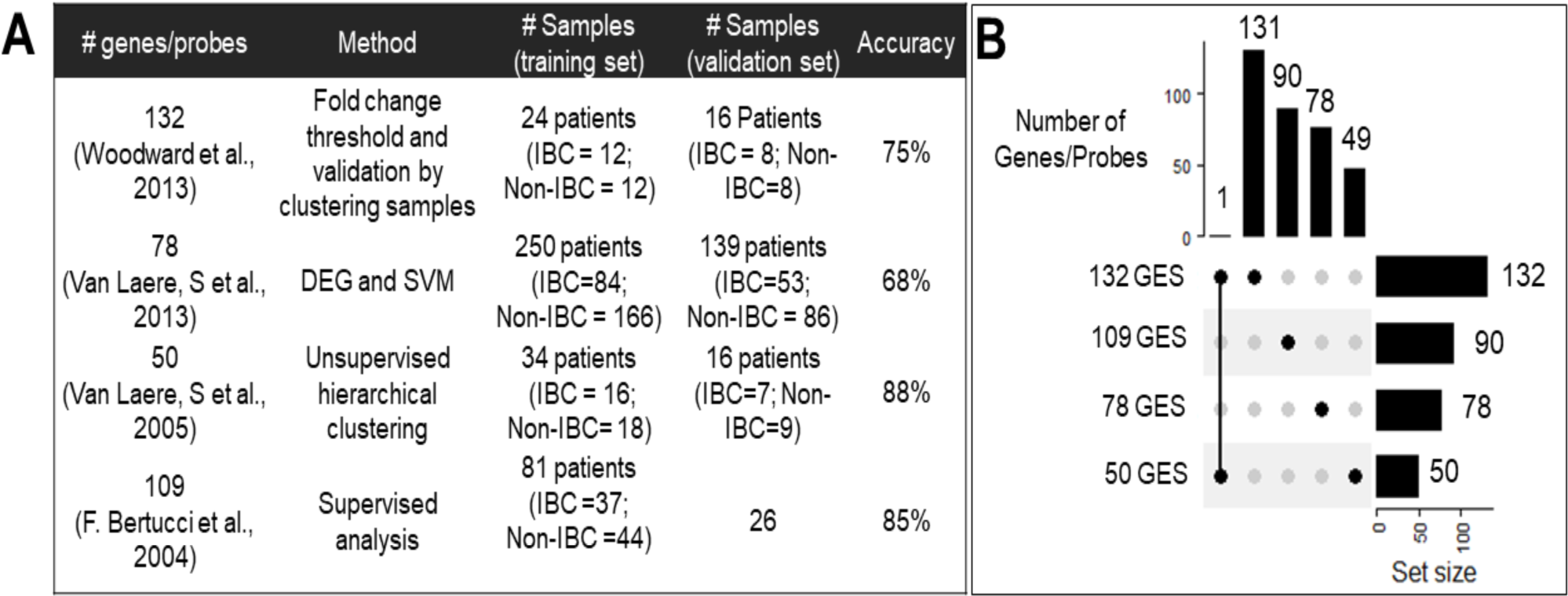
IBC gene signatures. **A)** Summary of all available IBC gene signatures **B)** Overlap in different IBC gene signatures

Based on high accuracy values (68% – 88%) of these gene lists in their ability to identify IBC in corresponding datasets, we hypothesized that each gene list might be capable of correctly classifying IBC and nIBC cases in three datasets containing gene expression data of clinically annotated IBC and nIBC samples (see Methods). We first performed principal component analysis (PCA) on all genes present in these datasets. In each case, PCA showed no clear separation between IBC and nIBC (**Fig 2A, i; Fig S1A i; S2A, i**), thus highlighting the high transcriptional heterogeneity observed in IBC and nIBC. Surprisingly, none of the four IBC gene lists (132 GES, 78 GES, 50 GES and 109 GES) performed consistently better than the all-genes approach in being able to segregate between IBC and non-IBC samples (**Fig 2A, ii-v; Fig S1A, B ii-iv**), with the only possible exception being the performance of 132 GES in one of the three datasets (**Fig 2A, iii**). This segregation was not improved by using non-linear methods such as uMAP as well (**Fig 2B, S1B, ii; S2B, ii**), suggesting that the mRNA levels of these genes are unable to resolve IBC and nIBC.

**Fig 2:**
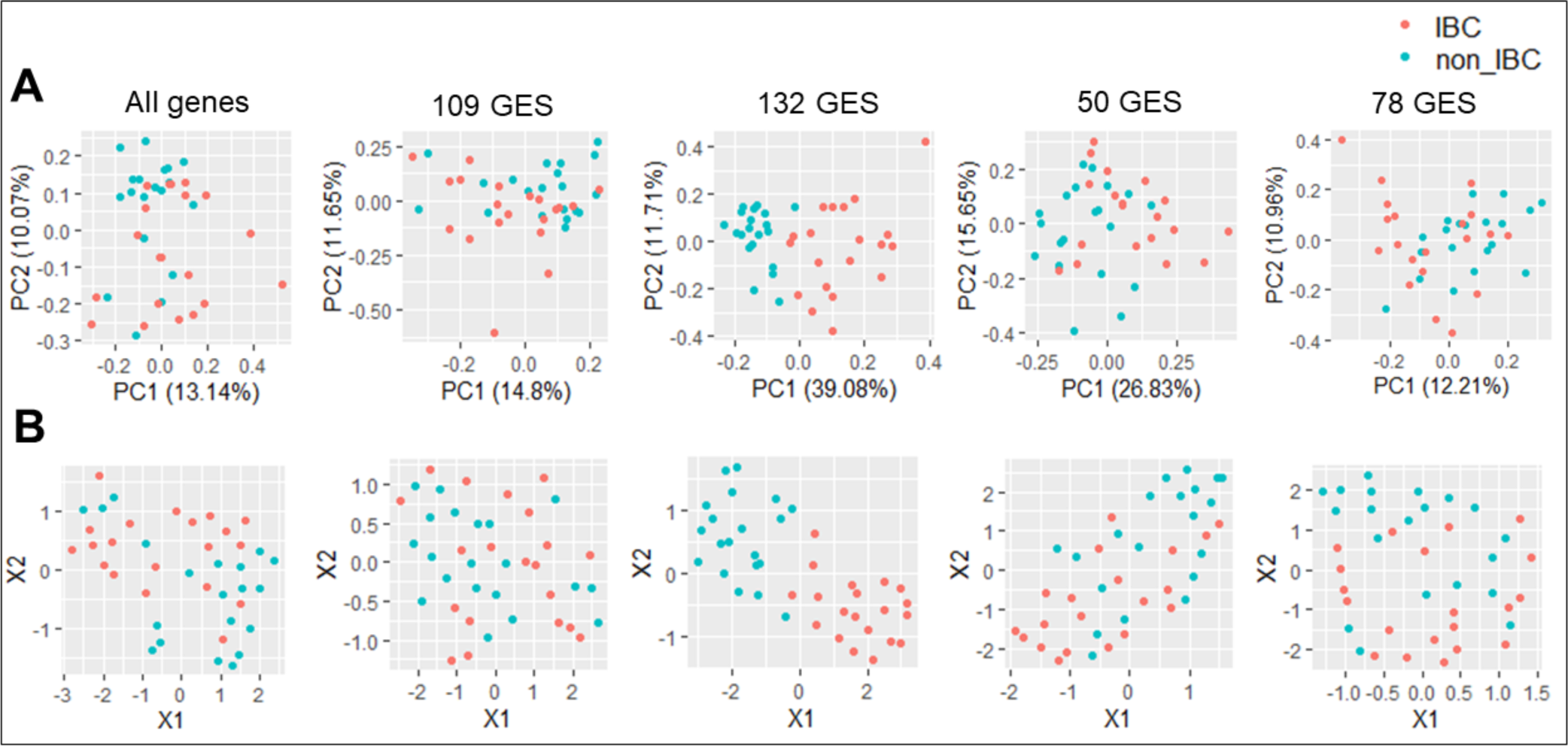
IBC gene signature expression in GSE45581 using dimension reduction methods. **A)** PCA **B)** uMAP

### ssGSEA scores of IBC gene signatures helps in separation of IBC and nIBC samples

As both linear and non-linear combinations of all four gene lists failed to segregate the IBC from nIBC samples in these datasets, we next examined whether as a group, these genes are enriched in IBC samples or not. We used single-sample GSEA (ssGSEA), an extension of Gene Set Enrichment Analysis (GSEA), which calculates separate enrichment scores for each pair of a sample and a gene set. Each ssGSEA enrichment score represents the degree to which the genes in a particular gene set are cumulatively up-or down-regulated within a given sample [14]. We thus tested whether IBC gene signatures are relatively enriched in IBC samples.

We calculated ssGSEA enrichment scores for all four different gene sets and compared these scores between IBC and nIBC samples across three corresponding datasets. This comparison showed some instances of statistically significant differences in ssGSEA scores of IBC and nIBC samples: a) ssGSEA scores of 78 GES for IBC samples was higher than that of nIBC samples in GSE22597, b) ssGSEA scores of 50 GES and 132 GES was relatively higher for IBC samples in GSE5847, and c) ssGSEA scores of 132 GES were comparatively higher for IBC samples in GSE45581 (**Fig 3A**). However, none of the four gene sets showed consistent and statistically significant differences between IBC and nIBC across the three datasets.

**Fig 3:**
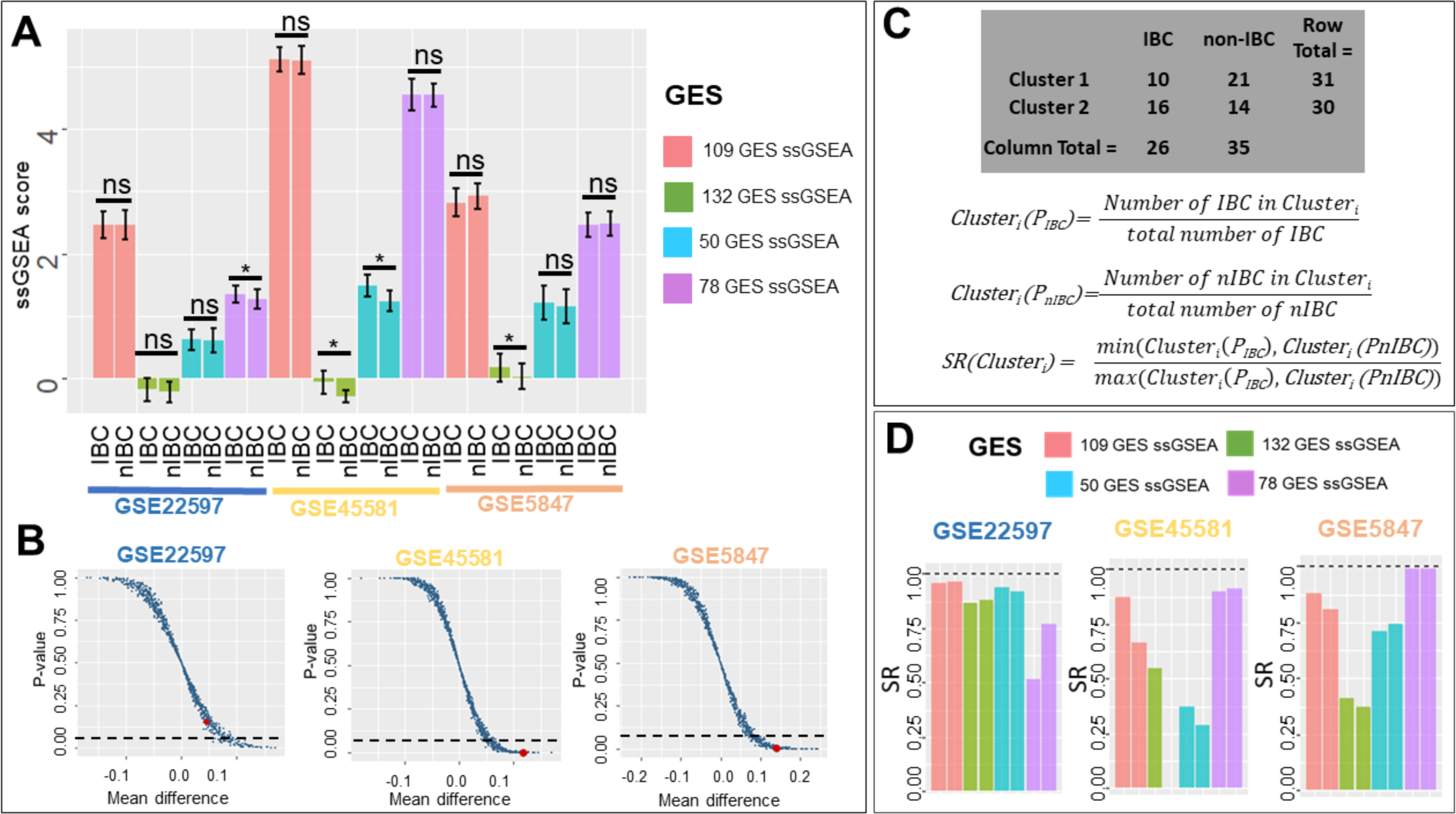
ssGSEA scores of IBC gene signatures. **A)** ssGSEA scores in IBC and nIBC samples across three datasets (* p< 0.05, two-tailed Student’s t-test; error bars represent standard deviation). **B)** Scatter plot showing ssGSEA scores calculated from 1000 random combinations of 132 genes each. Each data point in the plot represents data for a randomly generated gene list; the single point highlighted in red is calculated from 132 GES. x-axis is Mean difference = mean(ssGSEA_IBC_) – mean(ssGSEA_nIBC_) and y-axis is p-value estimated by *two-tailed Student’s t-test* of ssGSEA score across IBC and nIBC group (dotted lines denote p-value = 0.05). **C)** SR (sample ratio) method to measure accuracy of clustering based on ssGSEA scores (GSE5847 clusters) **D)** SR (sample ratio) of different gene signature ssGSEA scores across three datasets.

Next, we performed a permutation test to check whether the statistical differences observed in the mean ssGSEA scores of IBC and non-IBC samples were specific to these gene lists. This test was performed for 132 GES, because it showed significant results in terms of being enriched in IBC samples vs. non-IBC ones (p<0.05) for two out of three datasets. We chose the same number of genes (132) randomly out of the entire list of genes available to calculate the ssGSEA scores of IBC and non-IBC samples and tested whether the means of ssGSEA scores were significantly different for IBC and non-IBC samples. We repeatedly generated 1000 such instances and compared it to the ssGSEA scores obtained for 132 GES for each instance. This experiment showed that across the three datasets, for a large number of such randomly chosen gene lists, the difference in mean values of ssGSEA scores was not statistically significant (**Fig 3B**). This analysis indicated that the predefined 132 GES is quite likely to better distinguish between IBC and non-IBC compared to a randomly chosen gene set, but the extent of utility of this signature needs further validation.

After comparing ssGSEA scores, we next tested their ability to sort samples into two clusters and checked whether those two clusters corresponded to IBC and nIBC samples. We performed k-means (k=2) clustering on all four ssGSEA scores to cluster the samples and measured the accuracy of clustering into IBC/nIBC based on the SR (sample ratio) values (**Fig 3C**). A perfect clustering would mean that one of two clusters identified via k-means contains all IBC samples in that dataset, and the other contains all nIBC samples. To quantify the effectiveness of clustering into IBC/nIBC, we first calculated cluster IBC and nIBC scores for both clusters; these are the percentage of IBC (and nIBC) samples in the entire dataset that get classified into this cluster. The closer one of these percentages is to 100% and the closer the other percentage is to 0%, the more exclusive (or accurate) the clustering. Thus, we calculated the sample ratio (SR) by dividing the smaller of these two cluster scores by the larger of these scores. For a given cluster, the further the SR value is from 1 or the closer the SR value is to 0, the closer that cluster is to contain either all IBC or nIBC samples. For GSE22597, clusters formed on basis of 78GES had the lowest SR values compared to clusters formed by 132 GES, 50GES or 109 GES. For GSE45581, the SR values of 132 GES were shown to be the least; for GSE5847, 132 GES performed the best (**Fig 3C**).

We considered the actual sample groups and cluster numbers as two categorical variables. Statistical significance of enrichment of IBC and nIBC samples across these two clusters for each dataset was calculated based on a Fisher-exact test. 78 GES was the best performer in GSE22597, 50 GES in GSE45581, and 132 GES in GSE5847 (**Fig 3C-D**).

### Logistic Regression iteratively identifies an IBC signature

After exploring all the available IBC gene signatures, next we tried to define IBC signature by applying logistic regression (LR) to the three different datasets. LR is a machine learning-based classification scheme that can predict the probability of a given sample to belong to one of the two (or more) categories. The LR approach has been used to bin samples into epithelial, mesenchymal or hybrid epithelial/mesenchymal categories [15]. Applying LR to each dataset individually, all transcripts are ranked based on their ability to distinguish between IBC and non-IBC samples in the corresponding dataset. The generation of the specific IBC signatures yielded reasonable preliminary models of IBC vs. non-IBC (**Fig 4A**). Predictor goodness-of-fit and predictive accuracies correlated within each dataset, with the highest values observed in GSE45581 and the lowest in GSE22597 (**Table S1**). First, we identified which transcripts were best able to resolve IBC from non-IBC samples in individual datasets. While top transcripts varied across each dataset in their ability to separate IBC and non-IBC samples, with deviances ranging from 18.3 to 89.7 in the top ten from each dataset, their predictive accuracies were quite comparable – 75%-80% (**Table S1**).

**Fig 4:**
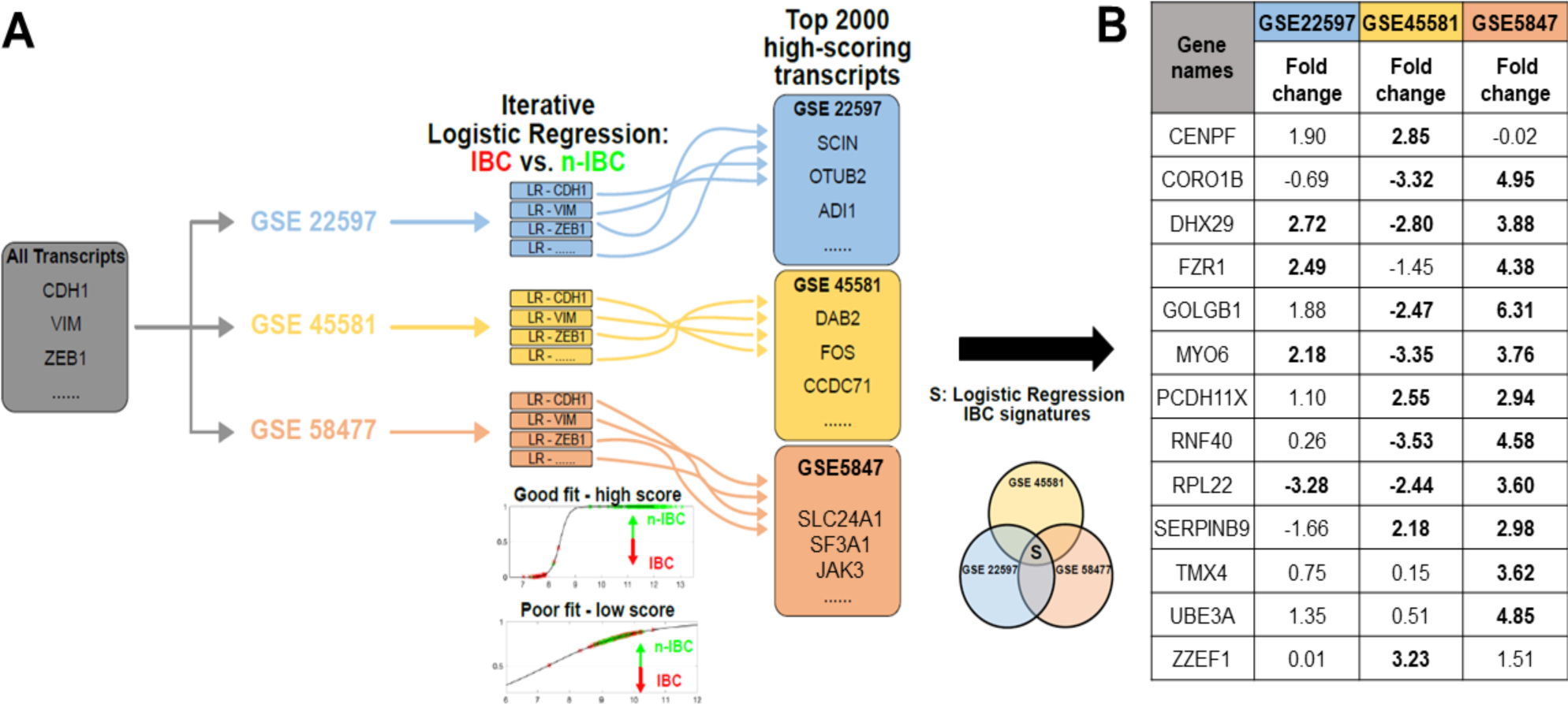
Identification of IBC-relevant Gene Signatures via Iterative Logistic Regression. **A)** Logistic regression was applied to each of the transcripts for GSE227597 (blue), GSE45581 (yellow), and GSE58477 (red). Transcripts were sorted based on their ability to resolve IBC and n-IBC samples. The intersection of the top 2000 transcripts were used to define the LR-specific IBC signature, S. **B)** LR signature fold-change between IBC vs. nIBC across three datasets. The bold-cases show when the fold-change value (mean (IBC)/mean (nIBC)) was statistically significant (p < 0.05) using a Students’ two-tailed t-test.

These results suggest that IBC can be reasonably identified, quite simply by using a single predictor, provided the analysis is restricted to a given dataset. On the other hand, generalizing across datasets is less straightforward. In an attempt to define an IBC signature based on the mutual intersection of three datasets, we looked for overlap in top ranked transcripts from each dataset; top 200 transcripts revealed no common genes. In top 2000 transcripts, 13 of them were found to be common across the three datasets. Third, we calculate the fold-change in levels of these 13 genes in IBC vs. non-IBC samples in these datasets. None of the 13 transcripts showed consistent upregulation or downregulation in IBC samples across the three datasets.

While our iterative approach represents the maximal resolvability achievable for each dataset using the iterative LR approach, the discrepancies across datasets seen in our earlier analysis do not completely disappear. For instance, lack of resolvability was also seen in IBC and non-IBC when using the LR-derived gene lists in PCA across the three datasets (**Fig S3 B-D**). Thus, increasing the predictive accuracy further may be a useful goal for future pursuits of a universal IBC signature. Such an approach is possible by utilizing multivariate LR models, but it requires significantly more training data for IBC and non-IBC cases, to prevent overfitting.

### Correlation between IBC gene signature ssGSEA scores and EMT scores

It has been proposed that IBC cells exhibit a partial EMT behavior, given the retention of E-cadherin levels and the trait of collective cell migration through tumor emboli [16]. Thus, following the assessment of IBC gene signatures, we quantified the EMT-ness of IBC and nIBC samples based on three different EMT scoring metrics – KS, 76GS and MLR [17].

These metrics score EMT on a continuum, based on the transcriptomics of individual samples. While KS and MLR score the samples on a scale of [-1, 1] and [0, 2] respectively, the 76GS metric has no pre-defined scale. The higher the MLR or KS score, the more mesenchymal the sample is; the higher the KS score, the more epithelial the sample is. Thus, KS and MLR scores of samples in a given dataset correlate positively with one another; both of them correlate negatively with 76GS scores, as seen across multiple datasets (**Table S2**)

Here, we used these metrics to estimate where IBC and nIBC samples lie in the entire epithelial-hybrid-mesenchymal spectrum, followed by an assessment of correlation of EMT scores with their corresponding ssGSEA enrichment scores. Two ssGSEA enrichment scores (132 GES and 50 GES) showed consistently very high and significant correlations with all three EMT scoring metrics in two datasets (GSE45581, GSE5847). Overall, a higher enrichment in IBC signature is associated with a more EMT-like phenotype. These correlations were maintained across both IBC and nIBC samples (**Fig 5A, B, Fig S6-S8**). However, this consistency is lost when using 78GES or 109GES in these datasets as well as in the case of GSE22597 using any of the four ssGSEA scores (132 GES, 50GES, 78 GES, 109 GES) (**Fig S3, S4**). Taken together, these results suggest that some of the IBC gene lists may enrich for mesenchymal samples instead of segregating IBC/ nIBC. The heterogeneity of both IBC and nIBC samples along epithelial-hybrid-mesenchymal spectrum may contribute to compromising the accuracy of these gene lists in identifying IBC.

**Fig 5:**
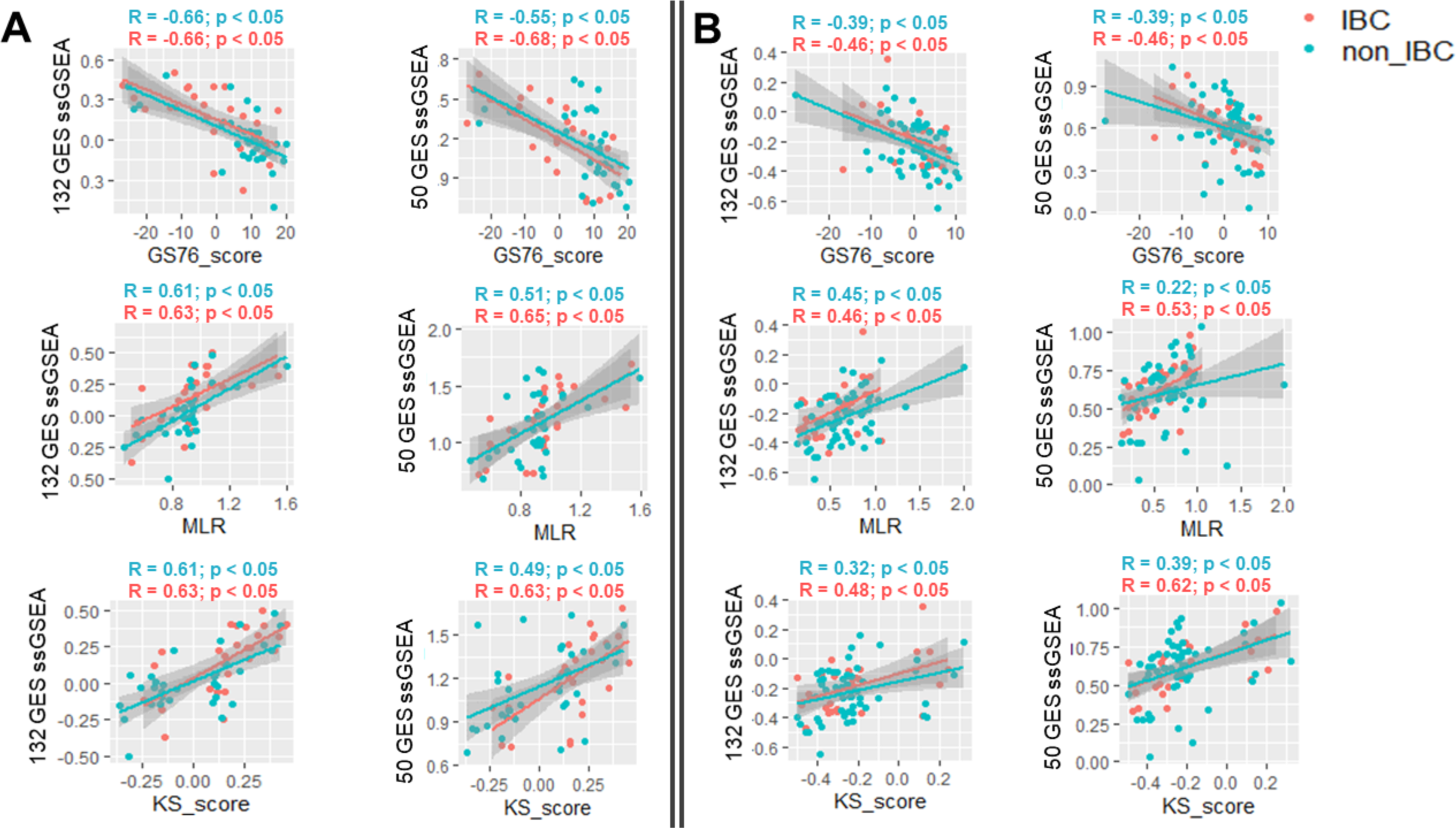
Correlation between ssGSEA score (132 GES and 50 GES) and EMT scoring methods. **A)** GSE5847 **B)** GSE22597. Pearson’s correlation R and p-values highlighted above each scatter plot

### Combination of EMT score and ssGSEA score helps in better separation of IBC and nIBC samples

Next, we asked whether IBC and nIBC can be separated using a combination of two different dimensions – ssGSEA enrichment score (using one or more of the four gene lists – 78 GES, 132 GES, 50 GES, 109 GES) and EMT score (using one or more of the EMT scoring metrics). To test the performance of different classifier combinations in sample segregation, we used different numbers and combinations of ssGSEA enrichment scores and EMT scores to perform k-means clustering. We used one, two, three, four, five, six, and seven different scores in all possible combination to cluster the samples into two groups. The clustering accuracy was again measured based on both SR value and a Fisher-exact test. This exercise showed that a combination of EMT scores and ssGSEA scores performs best in terms of clustering IBC and nIBC into separate groups (**Table S3**). Again, these combination of scores were not consistent across different datasets but it was always one or more ssGSEA scores with the combination of one or more EMT scores. Based on the SR value and Fisher exact test – (1) Combination of 78 GES and KS in GSE22597 (2) Combination of 50 GES, 132 GES, and KS in GSE45581 and (3) Combination of 132 GES, MLR, and KS in GSE5847 were the best performers in terms of clustering (**Fig 6A**).

**Fig 6:**
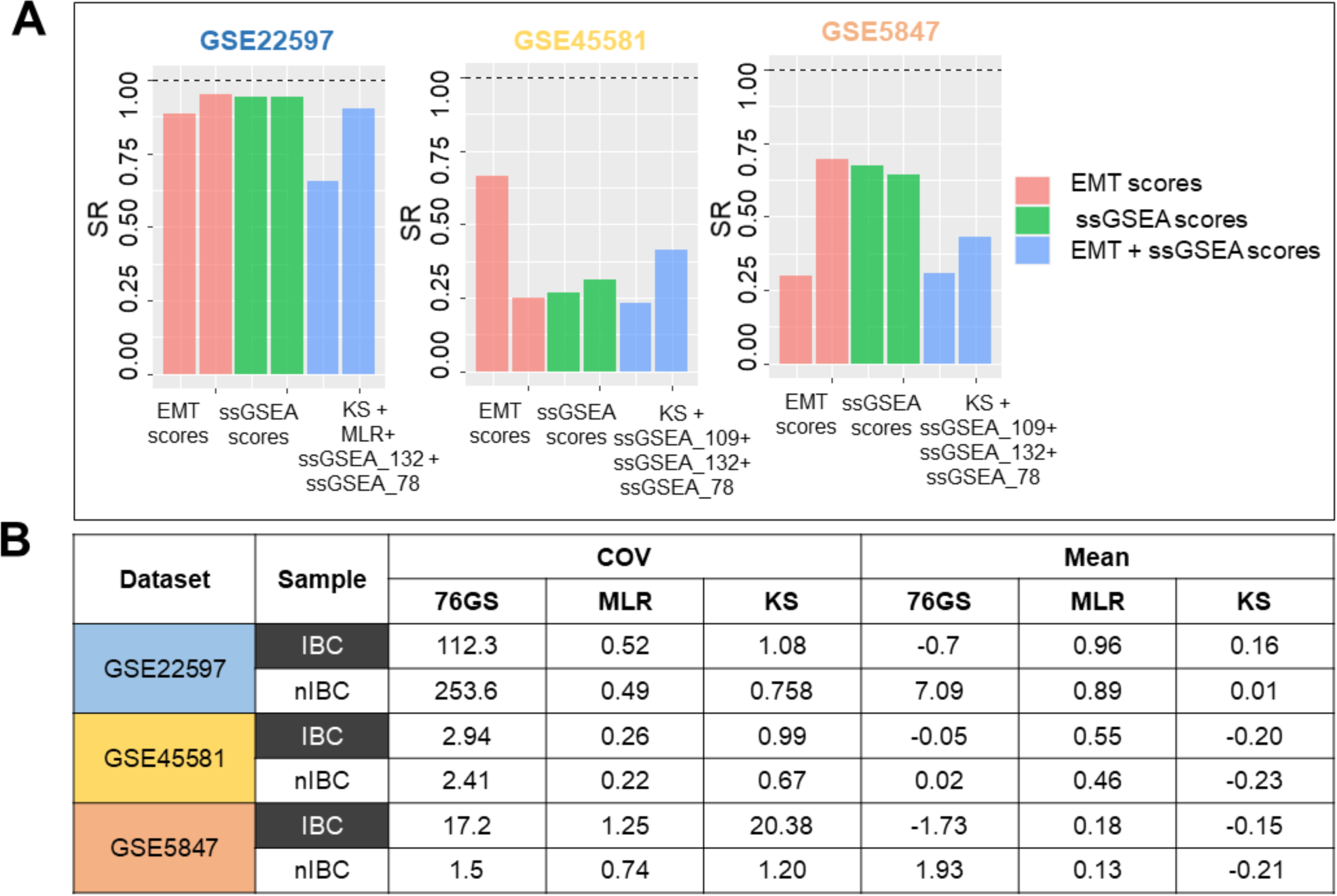
EMT scoring based identification of IBC. **A)** Clustering accuracy based on the combination of ssGSEA and EMT scores **B**) EMT Score mean and COV (coefficient of variation) across IBC and nIBC samples across the three independent datasets

### EMT score COV (coefficient of variance) is higher in IBC samples

Finally, after establishing the importance of EMT scores in segregation of IBC and nIBC samples, we compared these scores across IBC and nIBC. There was no significant difference in mean scores of IBC vs. nIBC across the three datasets (**Fig S9**), with overlapping variance between the two groups. However, the within-group coefficient of variance (COV), a better measure than variance to assess the dispersion around the mean, was consistently higher in 8 out of 9 (3 EMT scores x 3 datasets) total cases (**Fig 6B**). This result shows IBC samples are more heterogenous as compared to nIBC in terms of their positioning on EMT spectrum.

## Discussion

Identifying unique genomic or transcriptomic signatures for IBC has been a challenge, and this lack of consensus limits potential molecular therapeutic approaches to treat this rare but deadly disease [1]. The term ‘inflammatory’ for IBC originated from its physical appearance, which mimics an acute inflammation of the breast [18]. However, a useful association between the inflammatory phenotype and the cell’s omics has not yet been established.

An earlier study that attempted to characterize IBC based on clinical presentation as a distinct molecular entity concluded that “molecular subtype and inflammatory character are two independent features of breast cancers” [19], as the major molecular subtypes described for non-IBC were also found to exist within IBC. Nevertheless, there were multiple studies aimed at defining the molecular signature of IBC using genome-wide gene expression profiling. These studies used several different unsupervised and supervised methods to identify features related to IBC [6,10,12,13]. These gene lists have little to no overlap, and none of them so far have been interpretable as one or more common biological processes or pathways. The main objective of our study was not to predict the status of a new sample as IBC or non-IBC, but to validate the previously defined IBC gene signatures using available datasets consisting of IBC and non-IBC samples. These gene signatures were defined in individual datasets containing both IBC and non-IBC samples using specific statistical models, and we investigated if these gene lists are capable of identifying IBC and non-IBC in independent datasets. Here, we compare these previously defined gene signatures in their ability to classify IBCs from non-IBC, based on a analysis of three separate microarray datasets, none of which were directly involved in identifying the four gene lists (132 GES, 109 GES, 78 GES, 50 GES). Additionally, we also elucidate the EMT status of IBC samples using three different transcriptomics-based EMT scores. To the best of our understanding, this is the first study that contrasts different IBC gene signatures and EMT scoring across IBC datasets.

Mechanistic studies utilizing *in vitro* and *in vivo* models have revealed some markers for IBC, such as P-cadherin [20]. This molecule has also been proposed as a marker of the hybrid epithelial/mesenchymal (E/M) phenotype [21] due to its role in promoting collective cell migration and invasion [22,23] as well as tumor-initiating properties [24]; both of these properties are considered as hallmarks of hybrid E/M phenotype(s) [25]. P-cadherin (CDH3) is also a transcriptional target of NP63α [26], another potential ‘phenotypic stability factor’ (PSFs) for a hybrid E/M phenotype [16]. Similar to other PSFs [27–29], overexpression of P-cadherin associates with poor clinical outcome in invasive breast carcinomas [30]. Another pathway that has been reported to be enriched in IBC is the IL-6 pathway [16] which can promote Notch-JAG1 signaling [31]. JAG1 was reported as one of the top upregulated genes in collectively migrating cells [32] and its knockdown severely inhibited emboli formation in SUM149 IBC cells [31]. Moreover, given the role of IL-6 in mediating tumor-stroma crosstalk in IBC [33], it is possible that IL-6 mediates cell-cell communication both among IBC cells and with stroma. Despite these promising mechanistic insights, no accurate predictive signature for IBC exists.

Our results highlight that the proposed IBC gene lists so far (109 GES, 78 GES, 50 GES and 132 GES), despite showing a good accuracy in their corresponding validation datasets, show quite limited success in segregating IBC from non-IBC samples in independent cases. Various reasons may contribute to this result: inconsistency in the identification of IBC in different clinical samples, possible contamination by stromal cells in the samples investigated, and/or lack of metrics other than gene enrichment. To further indicate this failure of transferability, we used logistic regression (LR) methods to identify predictors (genes) that can best segregate between IBC and non-IBC, but no common trend was seen even in the top 2000 predictors collated from each of the three clinical datasets investigated. Put together, there exists a need to examine alternative metrics to be able to accurately identify IBC samples and perhaps gene expression on its own is insufficient to distinguish between IBC and non-IBC. One potential way to overcome the existing limitation may be to apply multivariate LR, but more training data to identify IBC from non-IBC samples would be required there to prevent any overfitting. Another approach would be to incorporate additional modalities of data, e.g. proteomics, to distinguish between IBC and nIBC.

Given the extensive literature on the role of a partial or complete EMT in collective migration and metastasis in breast cancer [32,34,35], we investigated if a more mesenchymal phenotype correlated with enrichment of ssGSEA scores for IBC gene lists in breast cancer samples. Indeed, ssGSEA scores for two IBC gene lists (50GES, 132 GES) correlated significantly with more mesenchymal samples, irrespective of whether those samples belonged to IBC or non-IBC. However, the ssGSEA scores for the 78GES list correlated with a more epithelial phenotype specifically for IBC samples, reminiscent of 78GES being associated with attenuated TGFβ signaling that may drive collective migration. Put together, one way to interpret these results may be that an ‘intermediate’ EMT associates with IBC, but at large, this inconsistency demonstrates a complex relation between IBC and EMT-ness and strengthens the idea of EMT-related heterogeneity in IBC. It is worth noting that these gene lists have been identified based on primary tumors and the expression signatures of primary tumors may contain little information about whether circulating tumor cells (CTCs) migrate individually or collectively [36]. Further characterization of emboli or clusters of CTCs for IBC will help deconvolute the contribution of EMT for IBC metastasis. Moreover, single-cell analysis of primary tumors and CTCs of IBC shall help in identifying various immune cell subsets in IBC which may drive disease progression. The composition and/or spatial localization of immune cells is likely to yield better insights into immune ecology of highly aggressive disease such as IBC [37,38].

The EMT scores for IBC as well as non-IBC samples did not indicate a ‘full-blown’ EMT (i.e. MLR score > 1.5 or equivalently KS score > 0.6), thus strengthening our previous observations that a complete EMT is not required for metastasis [39]. Intriguingly, we did observe a higher heterogeneity in EMT scores for IBC samples as compared to non-IBC samples. It remains to be ascertained whether the inter-tumor heterogeneity in EMT scores in IBC is also reflected as high intra-tumor heterogeneity as well. If that turns out to be the case, a higher intra-tumor heterogeneity along the EMT spectrum can be considered as a potential biomarker for IBC.

While genomic heterogeneity in tumors has been extensively studied, quantifying non-genetic (i.e. phenotypic) heterogeneity has been possible only recently through investigating cell-to-cell variability in isogenic populations [40–43]. Higher phenotypic heterogeneity may encourage cancer invasion [44] as well as the evolution of therapy resistance [45]. It may arise from network topology features such as mutually inhibitory feedback loops [46], for instance, the loop between RKIP and BACH1 [47] or that between AMPK and AKT [48]. Our earlier attempts have highlighted network topology based features such as hierarchical organization as a marker for IBC [39], endorsing that quantitative metrics to dissecting phenotypic heterogeneity in IBC may be better poised to highlight the IBC hallmarks than searching for gene signatures.

## Acknowledgements

This work was supported by Ramanujan Fellowship (SB/S2/RJN-049/2018) awarded to MKJ by SERB, DST, Government of India.

## Author contributions

PC and JTG performed research, HL and WW analyzed data, MKJ designed and supervised research. All authors participated in writing and editing of the manuscript.

## Methods

All the analyses have been performed on R 3.4.4 version and data was plotted using ggplot2 package.

### Datasets

Three separate IBC datasets were used in this study. GSE5847, GSE22597 and GSE45581 datasets were downloaded from NCBI GEO website.

### Principle component analysis (PCA)

PCA was performed using prcomp function available in R and plotted using factstoextra R package.

### ssGSEA analysis

ssGSEA analysis for various different gene sets were performed using GSVA R Bioconductor package with “ssgsea” option for method argument.

### k-means clustering

K-means clustering was performed using R package “cluster” and centers were set as two to get two separate clusters.

### EMT scoring

Three different EMT scoring methods – KS, MLR, 76GS were used to score samples separately in the three datasets [17].

### Statistical analysis

All the pairwise comparison significance was tested using student’s t-test. Significance of the enrichment of IBC and nIBC samples across clusters were tested using fisher exact test.

### Permutation/Randomization test

Permutation test was performed to test the significance of a gene signature as compared to the random set of genes. To determine the efficiency of an IBC gene signature on the basis of a distribution firstly the values were calculated using original expression values then same number of genes were chosen randomly. This process was repeated 1000 times to generate a null distribution of ssGSEA scores. Next, these ssGSEA scores compared across IBC and nIBC groups to obtain difference in mean values and significance based on t-test. This test was used to show that the original expression shows a higher difference in the mean of ssGSEA score and a lower p-value as compared to most of the random cases.

### Resolution IBC vs. n-IBC via iterative logistic regression

Samples from GSE22597 were first identified and categorized into IBC and n-IBC samples as previously reported, with all other samples omitted from analysis. The predictor set was comprised of all transcripts and for each individual transcript binomial logistic regression was fitted to the categorical IBC status. The output corresponds to each transcript a generalized residual sum of squared error, or deviance, with smaller values corresponding to better fit. The transcripts were then sorted in increasing order of deviance. For each of top ten predictors coding for genes, a leave-one-out assessment of prediction ability was performed. In each case, the logistic regression model was again constructed, this time on all but one sample. The corresponding statistical model was then used to predict the IBC status of the withheld sample. This procedure was repeated iteratively, withholding a distinct sample each time, and the results of the prediction were aggregated to estimate predictive accuracy.

This process was repeated for two additional datasets (GSE45581 and GSE58477). The top ten predictors for each dataset, together with their deviance values and predictive accuracy are listed in **Figure S3**. The top predictors from each of the three datasets define a logistic regression (LR)-specific IBC signature by taking the mutual intersection of the top 2000 scoring transcripts from each dataset (Figure 3).

## Supplementary figures

**Fig S1:**
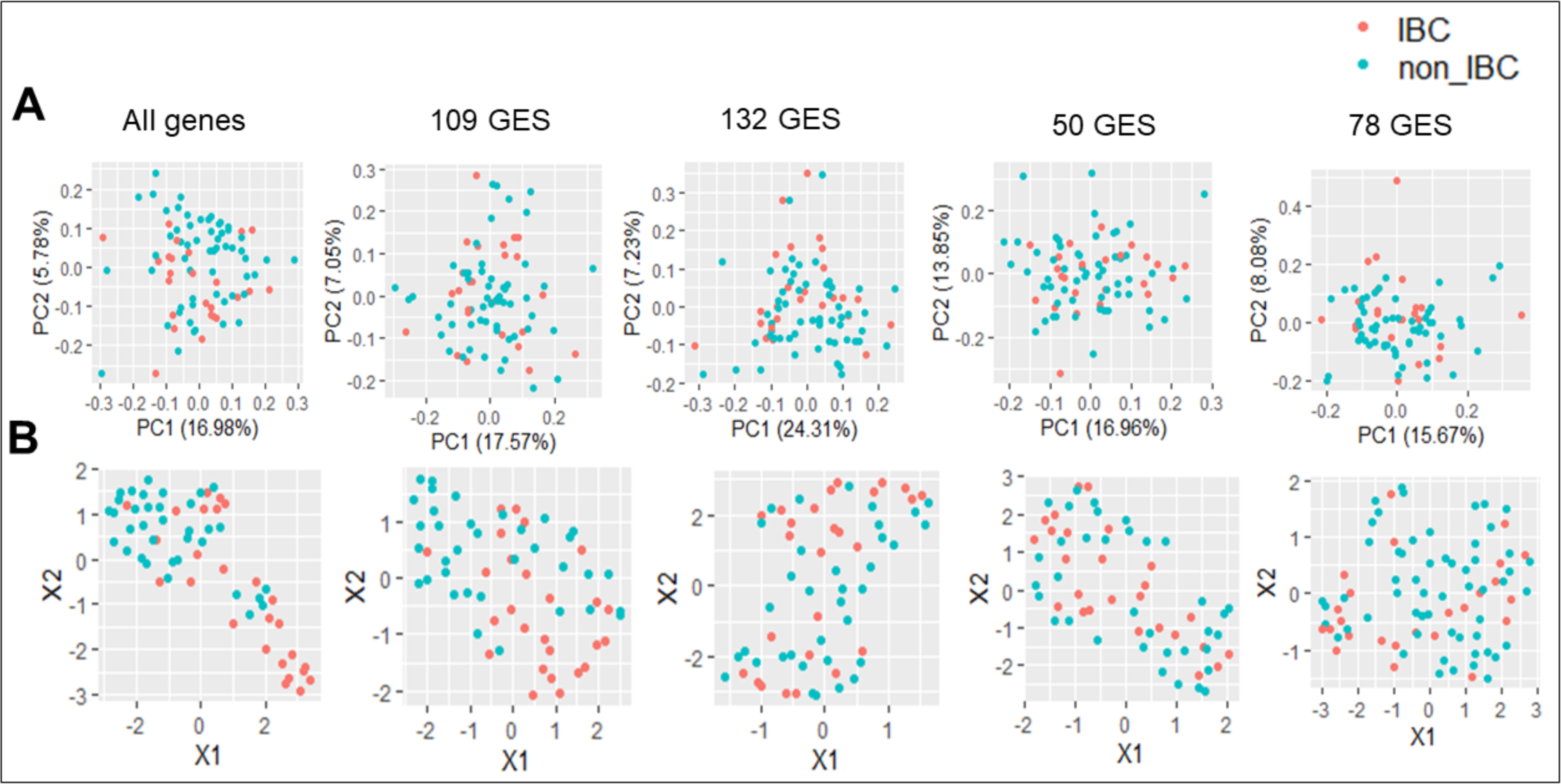
IBC gene signature expression in GSE22597. **A)** PCA **B)** uMAP

**Fig S2:**
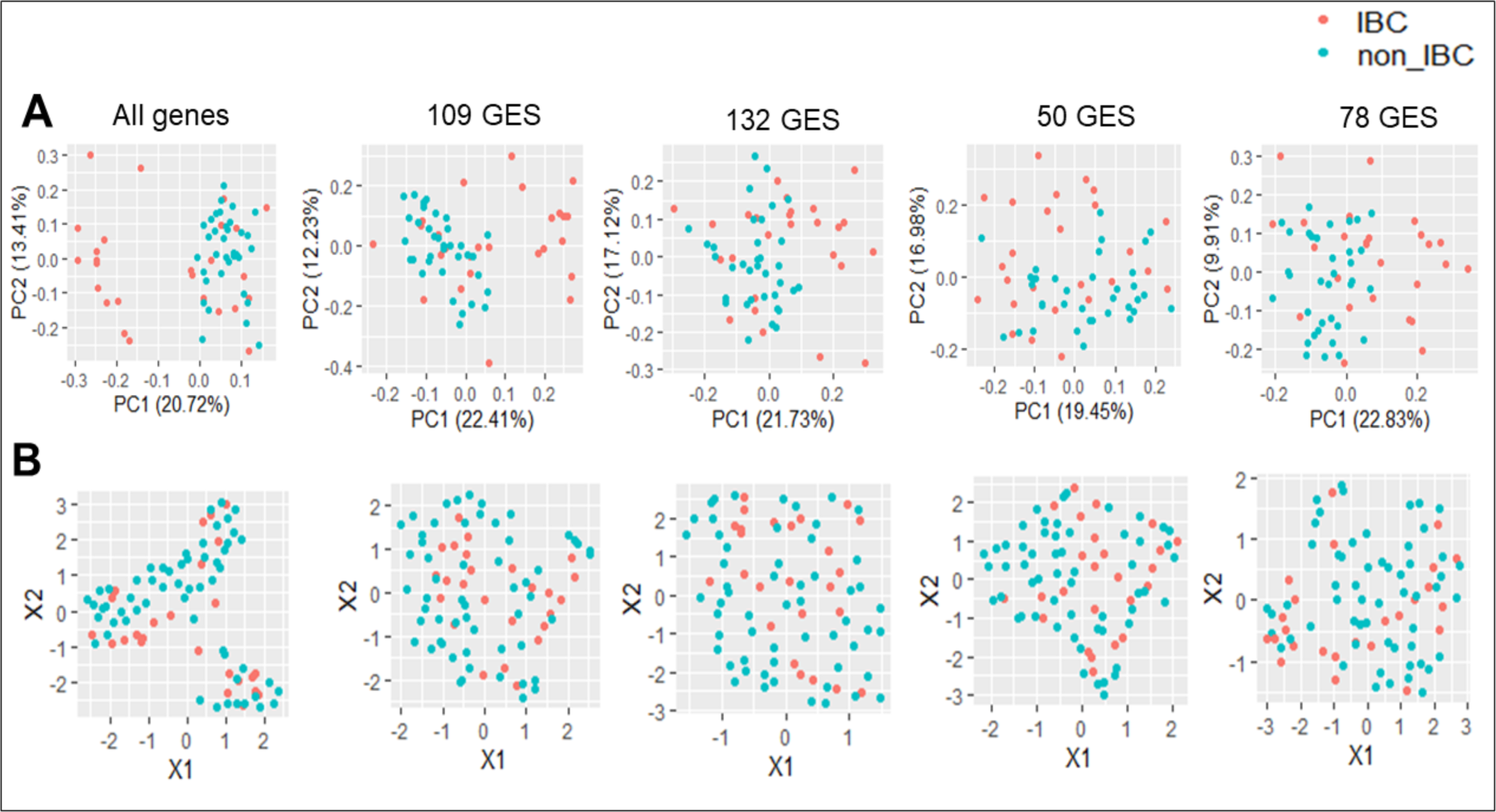
IBC gene signature expression in GSE5847. **A)** PCA **B)** uMAP

**Figure S3:**
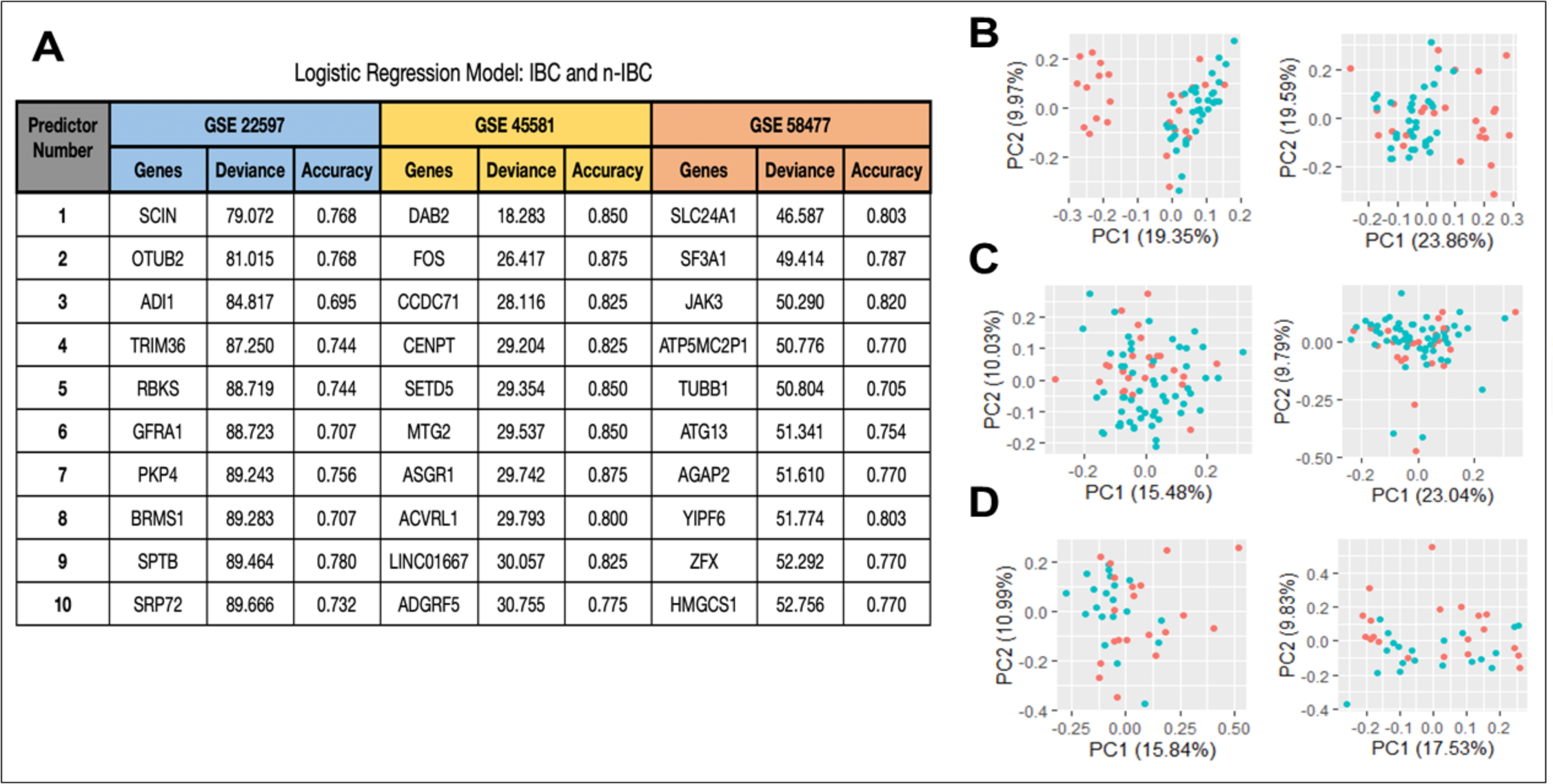
Features of the top LR predictors. **A)** The top 10 predictors for the GSE 22597 (blue), GSE 45581 (yellow), and GSE 58477 (red) datasets were identified based on their minimal deviance values. Leave-one out predictive accuracy is reported for each top transcript. **B)** GSE5847 PCA using top 100 LR predictors based on GSE22597 and GSE45581. **C)** GSE22597 PCA using top 100 LR predictors based on GSE5847 and GSE45581. **(D)** GSE45581 PCA using top 100 LR predictors based on GSE22597 and GSE5847.

**Fig S4:**
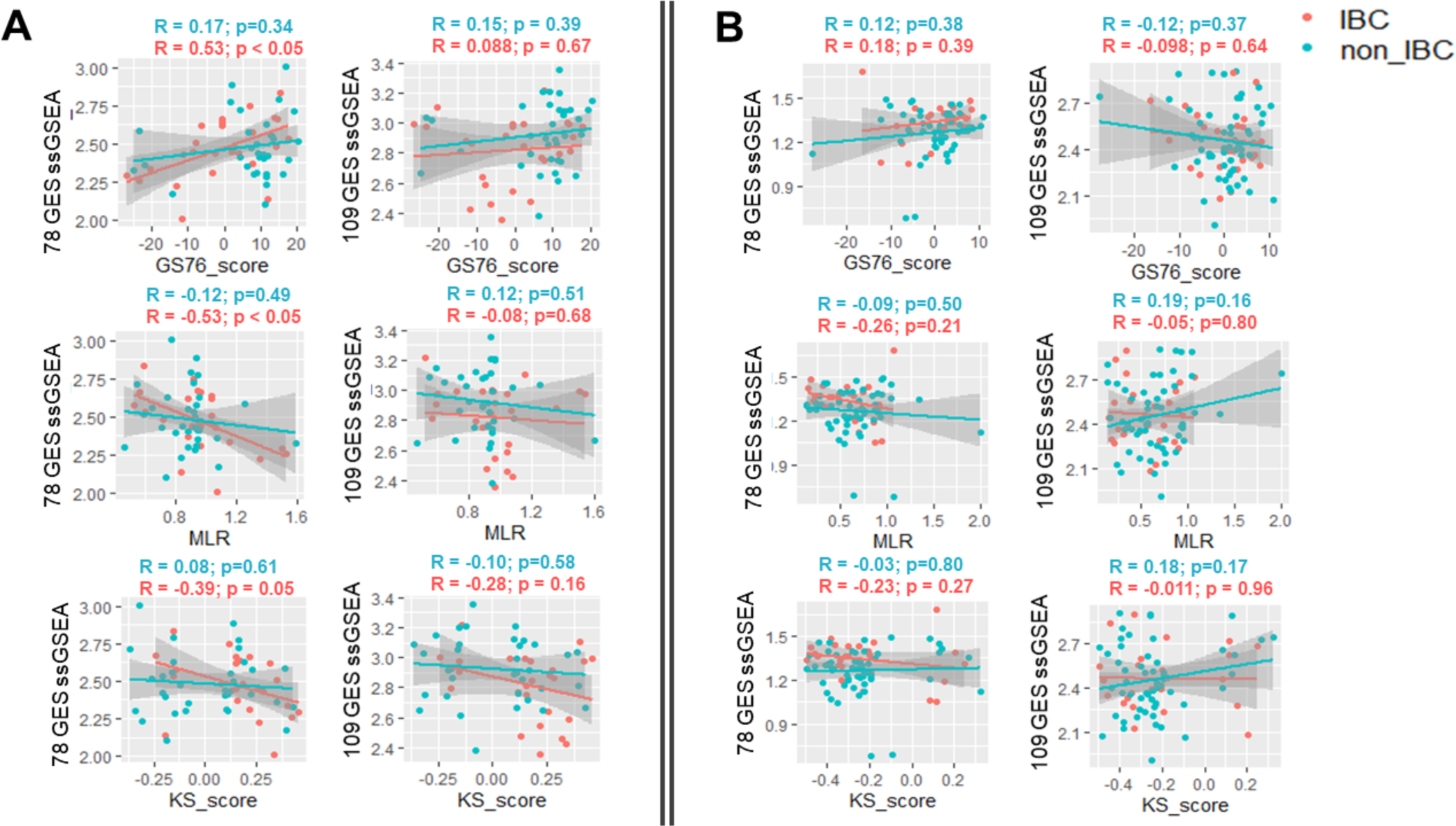
Correlation between ssGSEA score (78 GES and 109 GES) and EMT scoring methods. **A)** GSE5847 **B)** GSE22597. Pearson’s correlation R and p-values high-lighted above each scatter plot (blue – nIBC, red – IBC).

**Fig S5:**
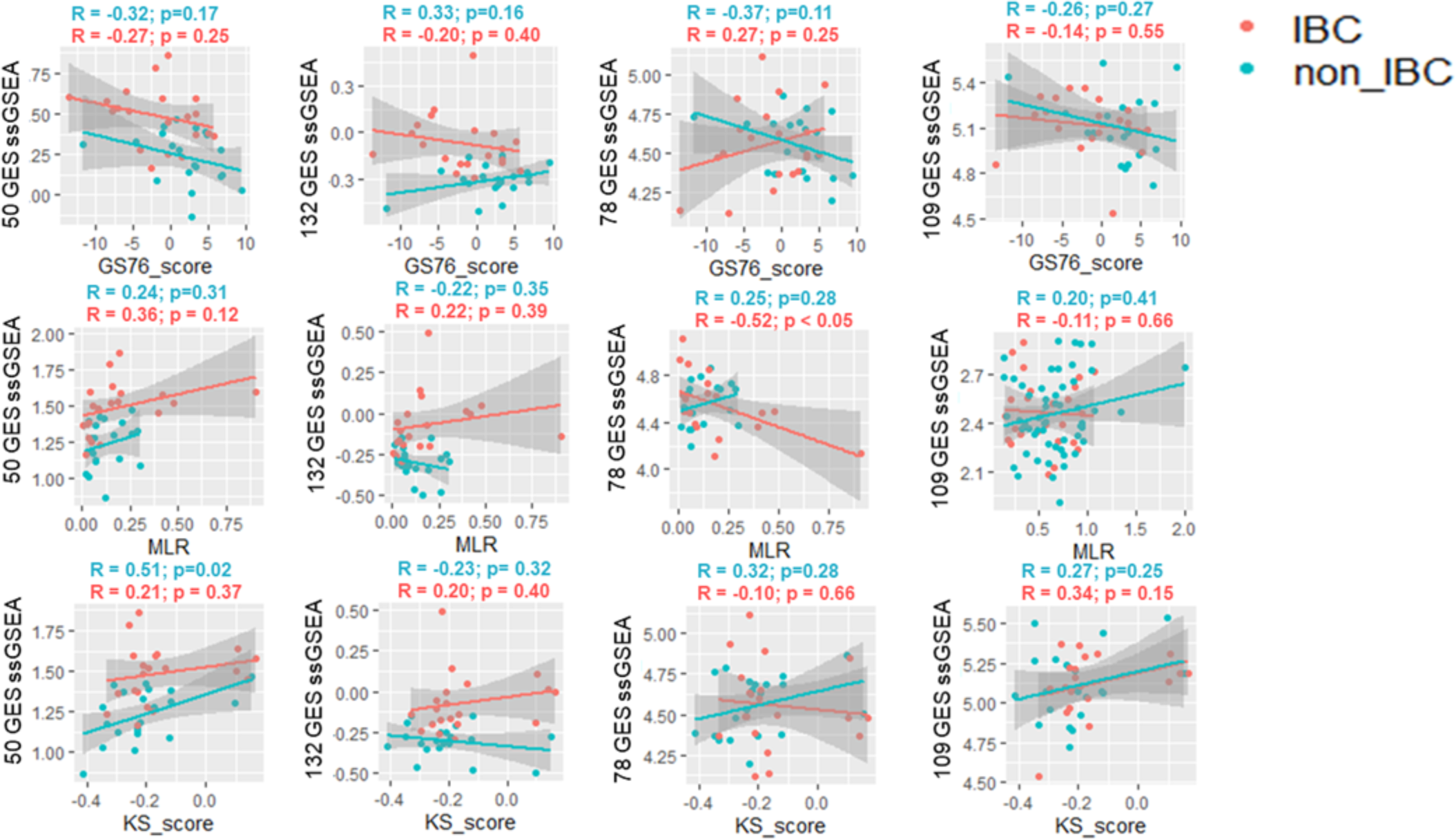
Correlation between ssGSEA score and EMT scoring methods in GSE45581. Pearson’s correlation R and p-values high-lighted above each scatter plot (blue – nIBC, red – IBC).

**Fig S6:**
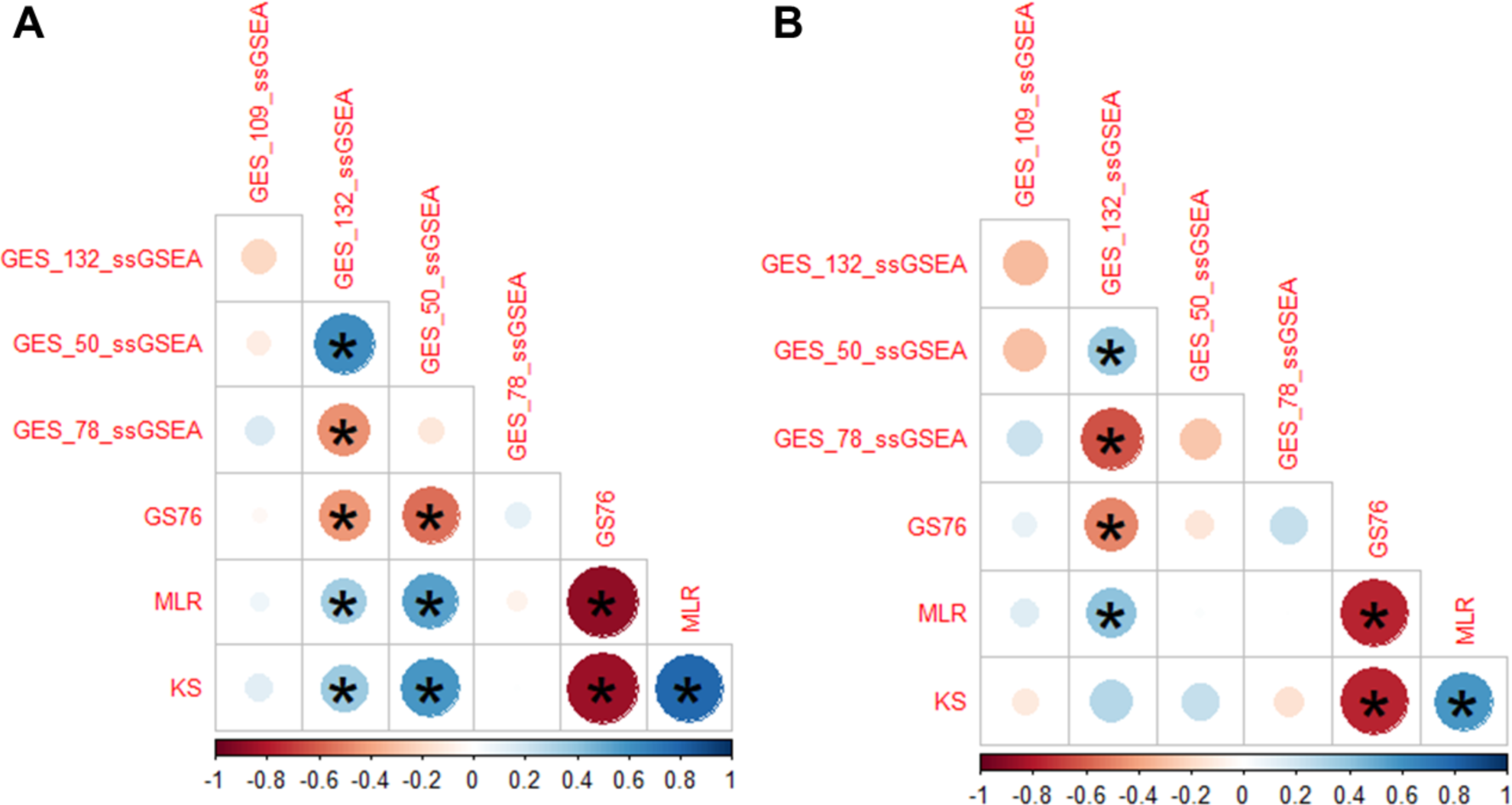
ssGSEA scores and EMT score Spearman’s correlation in GSE22597. **A)** IBC **B)** nIBC

**Fig S7:**
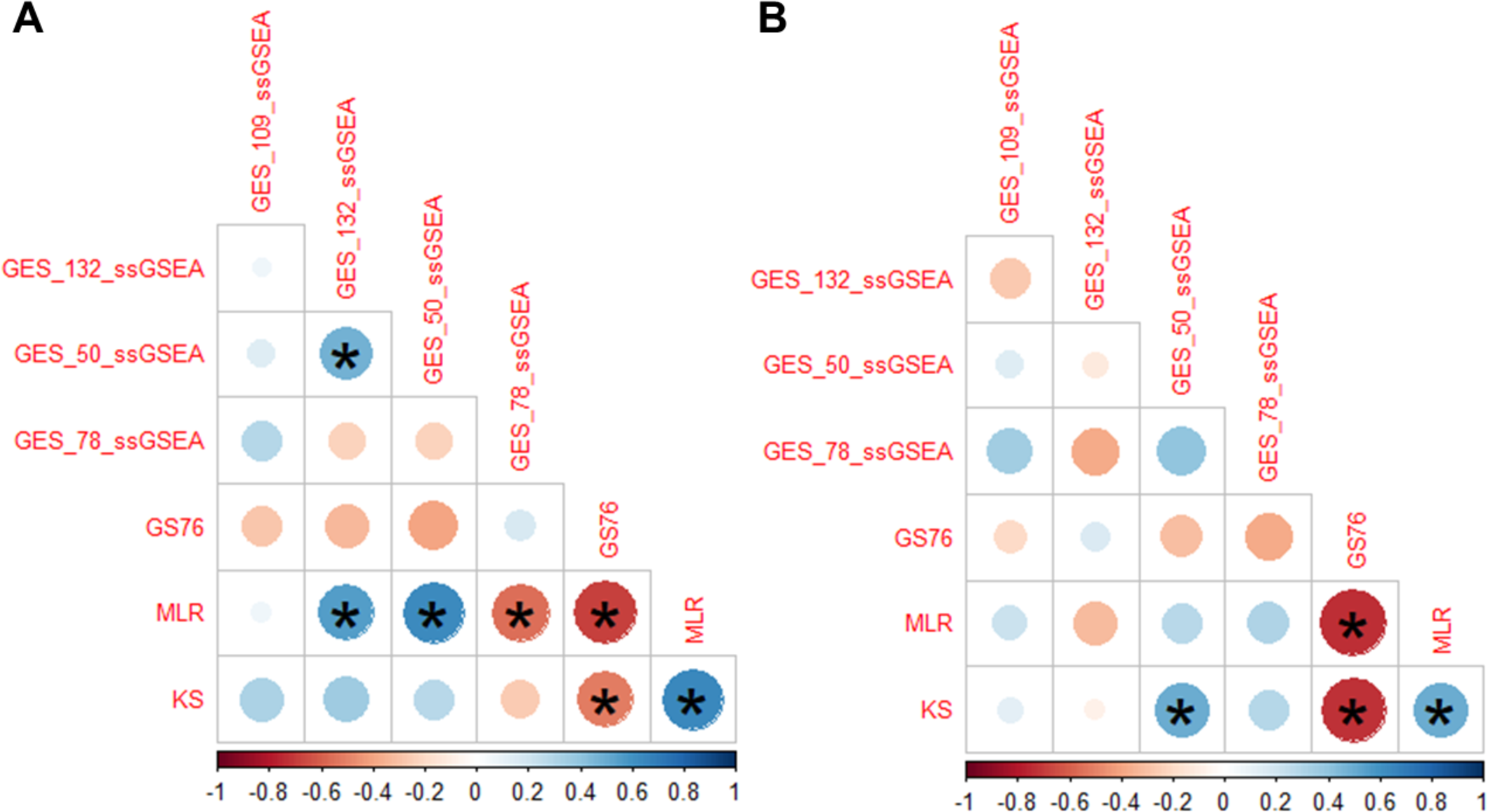
ssGSEA scores and EMT score Spearman’s correlation in GSE45581. **(A)** IBC **(C)** nIBC

**Fig S8:**
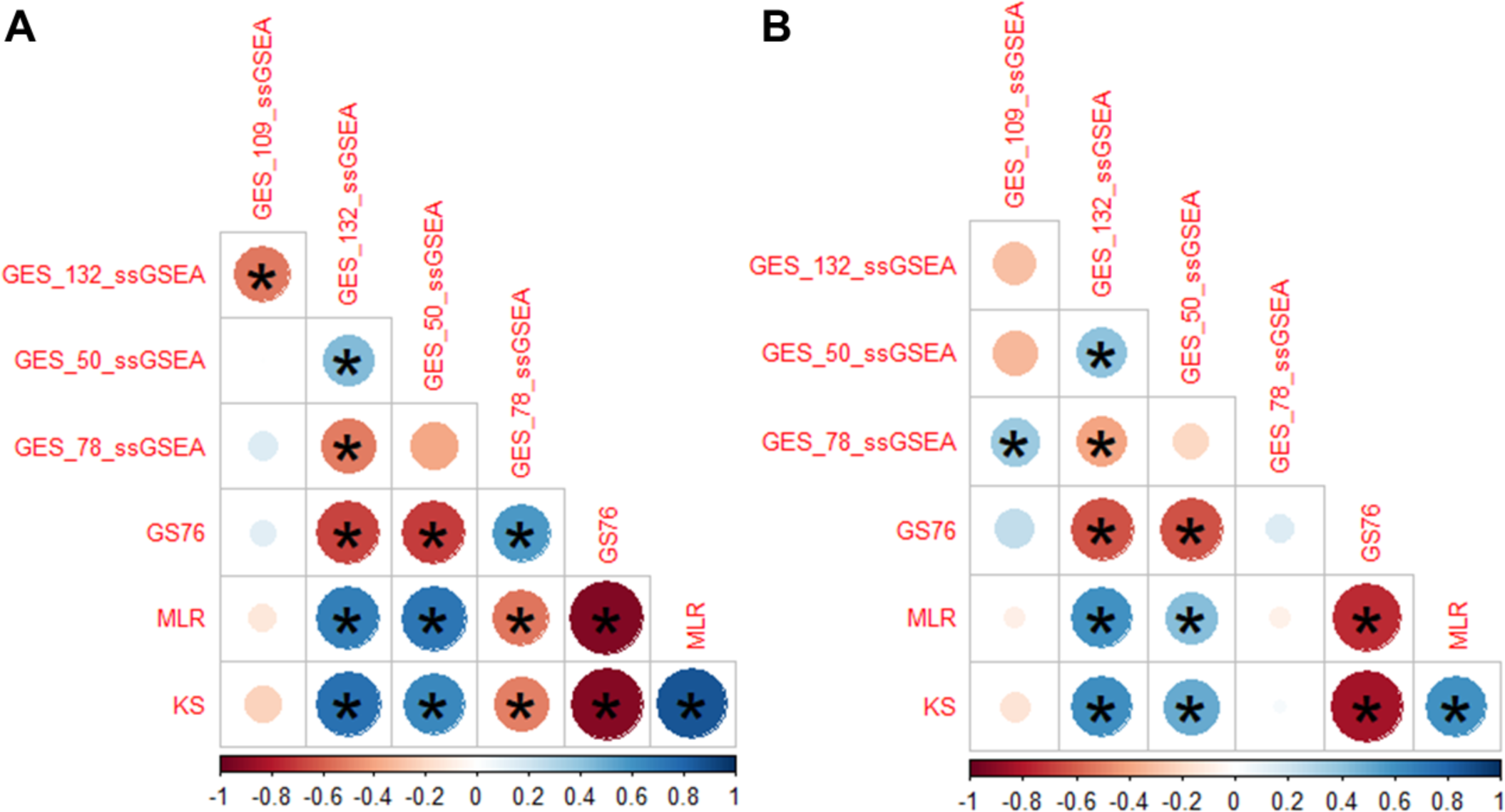
ssGSEA scores and EMT score Spearman’s correlation in GSE5847. **A)** IBC **B)** nIBC

**Fig S9:**
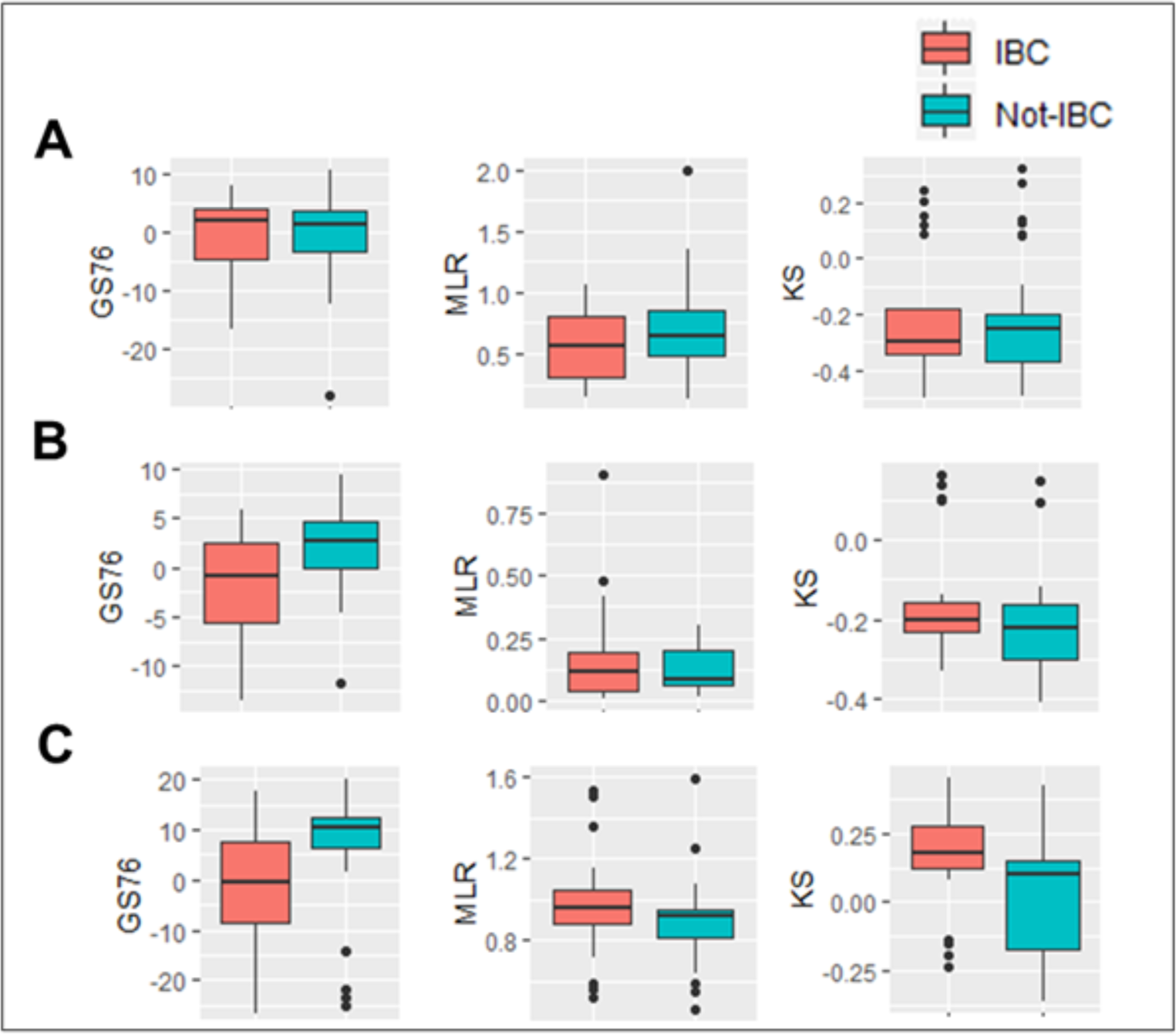
EMT scores across IBC and nIBC groups. **A)** GSE22597 **B)** GSE45581 **C)** GSE5847

## Supplementary table legends

**S1:** LR top 2000 predictors

**S2:** EMT scores

**S3:** Clustering accuracy of different combination of variables

